# Distinct CD8^+^ T Cell Programming in the Tumor Microenvironment Contributes to Sex Bias in Bladder Cancer Outcome

**DOI:** 10.1101/2020.04.13.039735

**Authors:** Hyunwoo Kwon, Dongjun Chung, Satoshi Kaneko, Anqi Li, Lei Zhou, Brian Riesenberg, No-Joon Song, Debasish Sundi, Xue Li, Zihai Li

**Author notes:** Corresponding authors: Correspondence to Zihai Li or Xue Li. **Disclosure of Potential Conflicts of Interest**: No potential conflicts of interest were disclosed.

## Abstract

Men and women show striking yet unexplained discrepancies in incidence, clinical presentation, and therapeutic response across different types of infectious/autoimmune diseases and malignancies^1,2^. For instance, bladder cancer shows a 4-fold male-biased incidence that persists after adjustment for known risk factors^3,4^. Here, we utilize murine bladder cancer models to establish that male-biased tumor burden is driven by sex differences in endogenous T cell immunity. Notably, sex differences exist in early fate decisions by intratumoral CD8^+^ T cells following their activation. While female CD8^+^ T cells retain their effector function, male counterparts readily adopt a Tcf1^low^Tim3^−^ progenitor state that becomes exhausted over tumor progression. Human cancers show an analogous male-biased frequency of exhausted CD8^+^ T cells. Mechanistically, we describe an opposing interplay between CD8^+^ T cell intrinsic androgen and type I interferon^5,6^ signaling in Tcf1/*Tcf7* regulation and formation of the progenitor exhausted T cell subset. Consistent with female-biased interferon response^7^, testosterone-dependent stimulation of Tcf1/*Tcf7* and resistance to interferon occurs to a greater magnitude in male CD8^+^ T cells. Male-biased predisposition for CD8^+^ T cell exhaustion suggests that spontaneous rejection of early immunogenic bladder tumors is less common in males and carries implications for therapeutic efficacy of immune checkpoint inhibitors^8,9^.

---

Sex is a biological variable with significant influence on immune function^1^. However, mechanisms underlying sex-biased incidence and mortality of various cancers arising in non-reproductive organs remain elusive^2^. Indeed, bladder cancer shows a 4-fold male-biased incidence globally, which cannot be explained by established risk factors: smoking, exposure to occupational hazards and urinary tract infection^3,4^. Here, we report that bladder cancer male bias, as replicated by multiple murine models, is largely mediated by sex differences in T cell immunity. Flow cytometric and single cell RNA-seq analyses of CD8^+^ tumor-infiltrating leukocytes (TILs) identified a striking male bias in the formation of Tcf1^low^Tim3^−^ progenitor cells^10^ and in their transition to a hypofunctional Tcf1^−^Tim3^+^ exhausted state upon prolonged stimulation. In particular, androgen signaling was enriched in the progenitor exhausted subset. We found that testosterone-induced Tcf1, an early molecular orchestrator of an exhaustion-associated transcriptional landscape^11^ in male CD8^+^ T cells, is also more resistant to pertinent repression by female-biased type I interferon signaling^5^. Collectively, these findings highlight sex differences in intratumoral CD8^+^ T cell fate as a potential mechanism underlying bladder cancer sex bias and identify androgen as a possible target to modulate CD8^+^ TIL exhaustion.

## CD8^+^ T cell immunity is required for sex differences in murine bladder cancer growth

Male bias in bladder cancer can be modeled successfully in mice with N-butyl-N-(4-hydroxybutyl)nitrosamine (BBN)-induced cancer^12,13^ and a transplantable syngeneic bladder cancer cell line MB49^14^. To assess the contribution of adaptive immunity to sex bias, we compared MB49 growth in male and female wild type (WT) mice, as well as mice with a deficiency of T cells (*Tcrb/Tcrd*^−/−^), B cells (*Ighm*^−/−^) or both (*Rag2*^−/−^). Consistent with an earlier report^14^, MB49 grew more aggressively in WT male versus female mice (**Fig. 1a**). However, sex differences in MB49 growth were eliminated in *Rag2* and *Tcrb/Tcrd,* but not *Ighm,* knockout (KO) mice, suggesting that they are driven by sexual dimorphism in endogenous anti-tumor T cell immunity (**Fig. 1a**). To confirm this finding, we repeated the experiment in WT mice after antibody-mediated depletion of CD4^+^ and/or CD8^+^ cells.

**Fig. 1.**
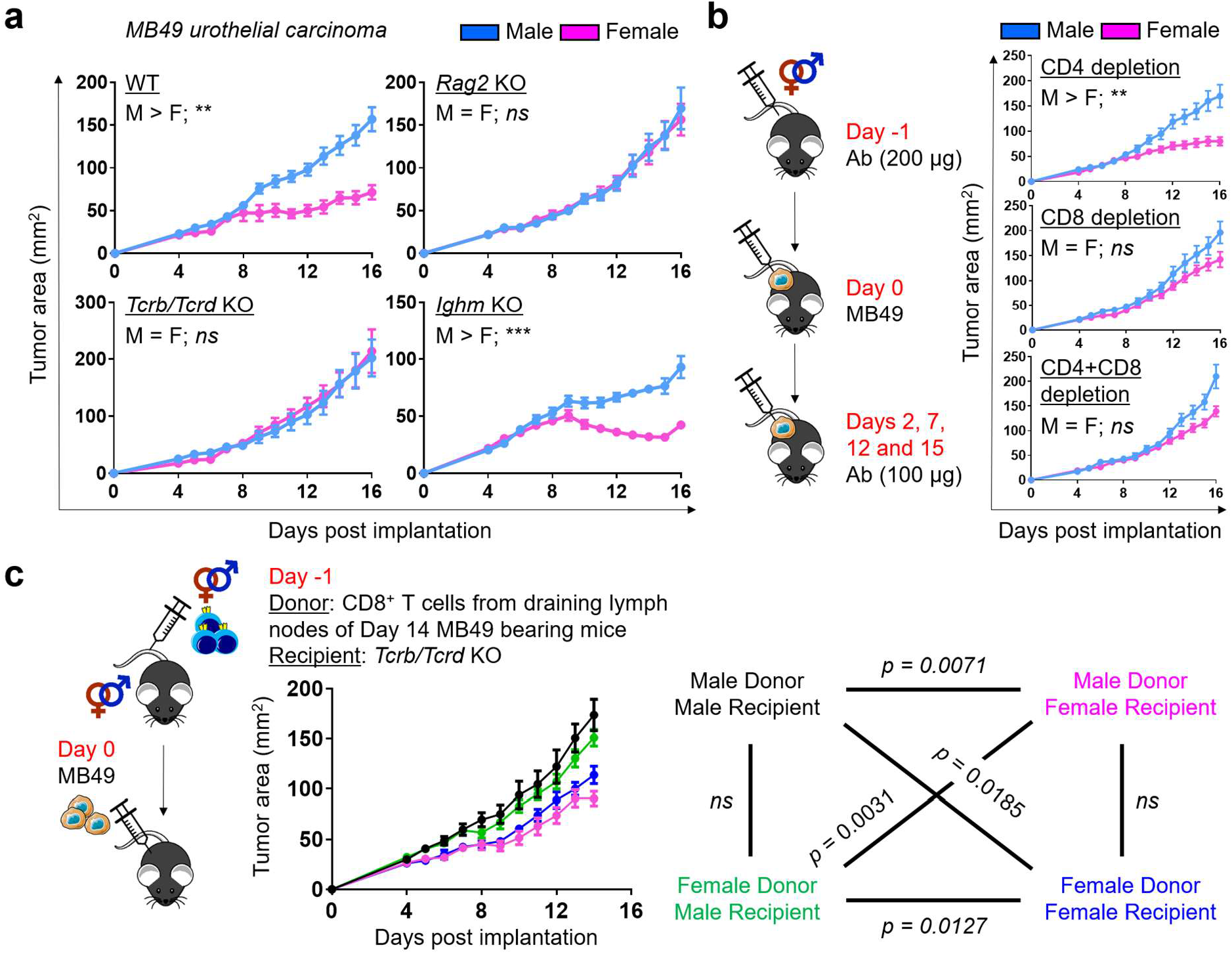
CD8^+^ T cell immunity is required for sex differences in murine bladder cancer growth. **a**, Growth of MB49 in mice with indicated genotypes after subcutaneous implantation. **b**, Antibody-mediated depletion of CD4^+^ and/or CD8^+^ cells in WT mice challenged with MB49. Each mouse was injected intraperitoneally with 200 μg anti-mouse CD4 (GK1.5 clone) and/or CD8 (53-6.7 clone) antibodies on day −1 and 100 μg on other indicated days. **c**, Growth of MB49 in *Tcrb/Tcrd* knockout mice that were adoptively transferred with 5 × 10^5^ CD8^+^ T cells from the draining lymph nodes of WT mice 14 days post subcutaneous MB49 challenge. Colors for all possible donor-recipient combinations and corresponding statistics are shown. **a-c**, Mean tumor area (mm^2^) ± SEM are reported, with statistical significance determined using the repeated measures two-way ANOVA. *n* = 4-10 mice per group. \**p* ≤ 0.05, \*\**p* ≤ 0.01, \*\*\**p* ≤ 0.001, *ns* = not significant.

We found that MB49 grew comparably in male and female mice specifically in the absence of CD8^+^ cells (**Fig. 1b**). Furthermore, we were able to induce sex bias in MB49 growth in *Tcrb/Tcrd* KO mice upon adoptive transfer of CD8^+^ T cells from the draining lymph nodes of MB49-bearing WT mice (**Fig. 1c**). Notably, the sex of recipients exerted a more significant influence on tumor suppression. These data indicate that CD8^+^ T cells, one of the primary mediators of anti-tumor immunity, are required for sex differences in the MB49 model.

While MB49 was generated through *in vitro* carcinogenesis of male mouse urothelial cells^15^, enhanced immune responses in female mice are not directed to male-specific minor antigens, as MB49 has lost the Y chromosome^16^, which was confirmed by a lack of *Sry* mRNA expression (**Extended Data Fig. 1a**). Furthermore, we generated a new syngeneic cell line “BKL171” from a BBN-induced bladder tumor in a testes-bearing Four Core Genotype (FCG) mouse with XX chromosome complement^17^ (**Extended Data Fig. 1b**). A clear male-biased tumor growth was also seen with BKL171 that was similarly dependent on T cell immunity (**Extended Data Fig. 1c**). Finally, we examined sex bias with the BBN-induced orthotopic bladder cancer model. Consistent with earlier reports^12,13^, WT female mice survived longer after BBN exposure. However, *Tcrb/Tcrd* KO mice lacked sex differences in BBN-induced carcinogenesis (**Extended Data Fig. 1d**). Collectively, our findings with three murine bladder cancer models – MB49, BKL171 and BBN-induced carcinoma – highlight a previously underappreciated role for sex differences in T cell immunity, especially CD8^+^ T cells, which drive male-biased bladder tumor growth.

## Female bias of CD8^+^ T cell effector response in the tumor microenvironment

We hypothesized that fundamental sex differences exist in the behaviors of CD8^+^ TILs. Recognizing that immune phenotypes often correlate with the magnitude of tumor burden, we focused our analysis on days 9 to 11 post MB49 tumor implantation, which coincides with the time of bifurcation in tumor growth curves for males and females, to identify a sex-specific driver – not passenger – phenotype. We observed that the number of MB49 tumor-infiltrating and peripheral CD45^+^, CD4^+^ and CD8^+^ T cells are comparable between sexes (**Extended Data Fig. 2a–2g**). However, there was an approximately two-fold higher frequency of polyfunctional CD8^+^ T cells that could produce Interferon gamma (IFNγ), Tumor Necrosis Factor Alpha (TNFα) and Granzyme B (Gzmb) in day 9 MB49 tumors, but not in spleens, of female versus male mice (**Fig. 2a**). A similar female bias in effector response was seen with tumor infiltrating CD4^+^ T cells, albeit at a lower magnitude (**Extended Data Fig. 3a–3b**). Next, we performed single-cell RNA sequencing (scRNA-seq) on 26,698 CD8^+^ T cells from day 10 MB49 tumors (9,955 from female and 16,743 from male; *n* = 3 mice per sex) to identify a molecular basis for observed sex-specific differences. The shared nearest neighbor modularity optimization-based clustering algorithm identified 11 major clusters of cells at various stages of T cell differentiation (**Fig. 2b and 2d; Supplementary Data 1**). Consistent with the immunogenic nature of MB49, all but cell clusters 2 and 7 showed signs of activation via T cell receptor (TCR), as marked by the expression of co-stimulatory receptors (*Cd28*, *Icos*, *Il2ra*), inhibitory surface receptors (*Pdcd1*, *Havcr2*, *Lag3*, *Ctla4*) and effector molecules (*Ifng*, *Gzma*, *Gzmb*). Clusters 2 and 7 were instead enriched for *Tcf7*, *Sell* and Ribosomes, which are characteristic of “stem-like”, “memory-precursor-like” or progenitor populations^5,10^. Additionally, clusters 3/5 and 9 were distinguished by the expression of genes associated with active cell proliferation (*Top2a*, *Mki67*, Histones, Tubulins and MCM complex) and type I Interferon signaling, respectively. *Tox* was not enriched in any clusters, indicating that none of the analyzed CD8^+^ T cells in day 10 MB49 had yet reached terminal differentiation^18–22^.

**Fig. 2.**
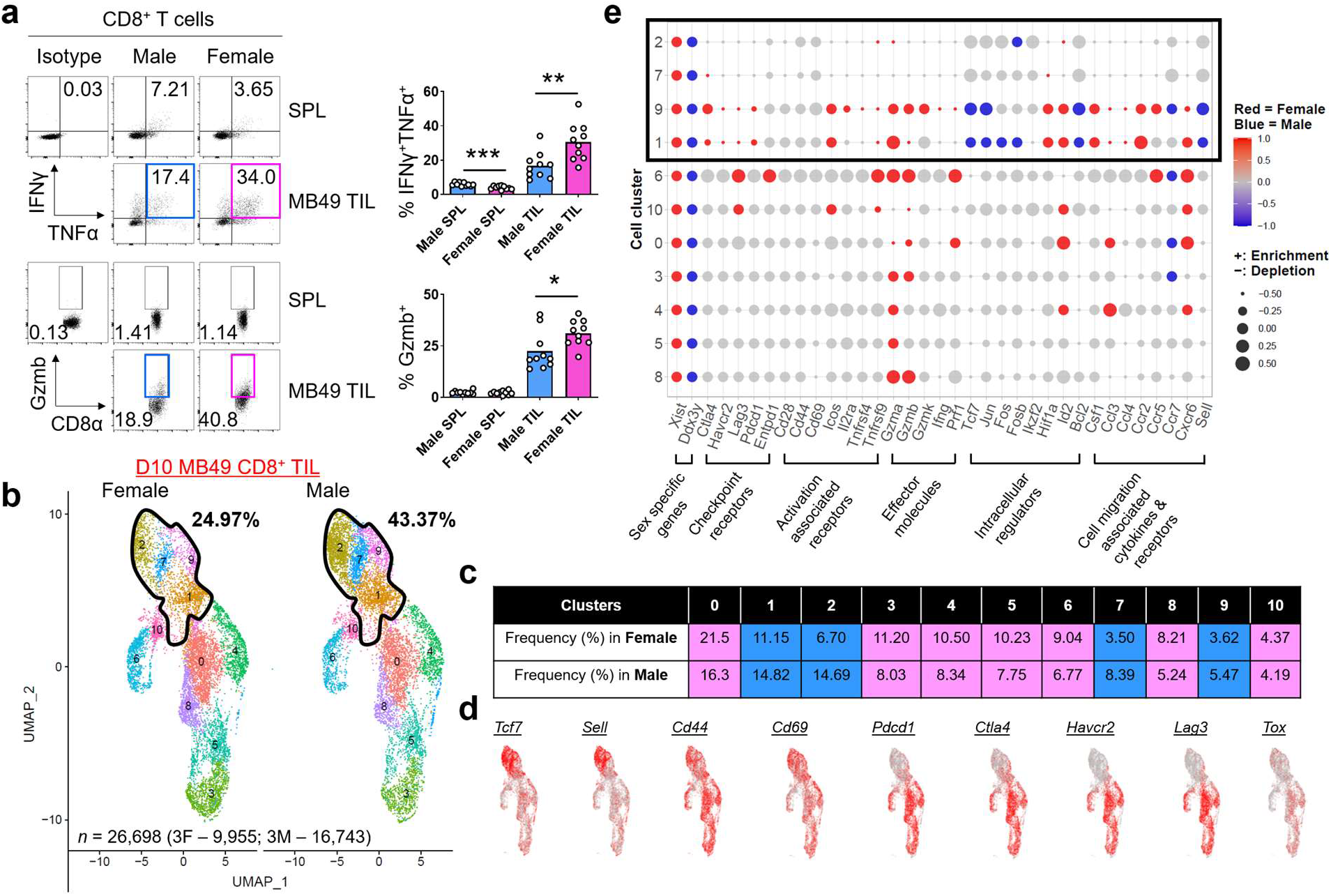
Female bias exists in CD8^+^ T cell effector response in the tumor microenvironment. **a**, Flow cytometric analysis of IFNγ and TNFα (top) and Gzmb (bottom) expression in CD8^+^ T cells from the spleens (SPL) and tumors (TIL) of male and female mice 9 days post subcutaneous MB49 challenge. Cells were stimulated *ex vivo* with 50 ng/mL PMA, 1 μg/mL Ionomycin and 1X Brefeldin A for 2 hours. Left: Representative flow plots; Right: Frequency of CD8^+^ T cells expressing indicated effector molecules, with the box height representing the mean of all shown biological replicates. Statistical significance was determined using the Student’s *t* test. \**p* ≤ 0.05, \*\**p* ≤ 0.01, \*\*\**p* ≤ 0.001. **b**, UMAP of 26,698 CD8^+^ TIL (3 female mice – 9,955; 3 male mice – 16,743) scRNA-seq profiles, colored by cluster. Cells were sorted by FACS from day 10 MB49 tumors. Same number of cells (9,955) is shown here for each sex for visualization. Solid black lines in **b** and **e** enclose 4 clusters that show male-biased frequency as indicated in blue in **c**. **d,** Expression of indicated genes in individual cells from **b**. **e**, Dot plot indicating the relative expression of a given gene in the 11 clusters and its sex bias by size and color, respectively. Sex-biased expression was assessed using a Mann-Whitney U test and its significance was determined at the nominal level of 0.05 after Benjamini-Hochberg multiple testing correction. Blue: Male bias; Red: Female bias; Gray: No sex bias.

We focused on identifying sex differences in the cluster frequencies and intra-cluster gene expressions (**Fig. 2c and 2e**). CD8^+^ T cells from clusters 2, 7, 1 and 9 were more predominantly found in MB49 tumors from male mice, with the first two demonstrating over two-fold higher frequency. All other seven clusters showed a female-biased frequency with increased transcripts for effector molecules. Clusters 1 and 9 appeared to represent an inflection, at which male and female CD8^+^ T cells diverge to adopt a stem-like versus effector fate, respectively. Specifically, clusters 1 and 9 were enriched for *Sell*, *Bcl2*, *Tcf7*, *Jun*, *Fos* and *Fosb* transcripts in males. By comparison, these clusters were enriched for co-stimulatory receptors (*Icos* and *Tnfrsf9*), inhibitory surface receptors (*Pdcd1*, *Havcr2*, *Lag3* and *Ctla4*), effector molecules (*Gzma* and *Gzmb*), transcription factors (*Hif1a* and *Id2*), chemokines/cytokines (*Ccl3*, *Ccl4* and *Csf1*) and migratory receptors (*Ccr2* and *Cxcr6*) in females. In conjunction with flow cytometric analyses (**Fig. 2a**), we conclude that the female tumor microenvironment favors development of effector-like CD8^+^ T cells with an enhanced ability to control tumor progression.

## Male bias exists in CD8^+^ T cell commitment to exhaustion in the tumor microenvironment

T cell exhaustion defines a state of dysfunction that arises upon persistent antigen stimulation such as in cancer or chronic infection, with progressive loss of effector and proliferative potential, sustained expression of inhibitory receptors and a distinct transcriptional landscape^23^. Progenitor exhausted CD8^+^ TILs, as defined by their stem-like genetic profile (*i.e.* Tcf1/*Tcf7*) with relatively little to no expression of checkpoint receptors, have the potential to proliferate and give rise to effector-like cells, particularly in response to anti-PD1 therapy^10,24–30^. Given that Tcf1/*Tcf7* plays a critical role in orchestrating a progenitor exhausted fate while antagonizing an effector program at the early stage of CD8^+^ T cell differentiation^11^, we postulated that male bias in *Tcf7* gene expression for clusters 1 and 9, as well as in the frequency of *Tcf7*-enriched clusters 2 and 7, may indicate development of progenitor exhausted CD8^+^ TILs. To infer their ontogeny, we used the Monocle-2 algorithm to order single cells in clusters 1, 2, 6, 7, 9 and 10 in pseudotime^31^. This analysis revealed a trajectory originating at State 1, which then bifurcated into States 2 and 3 (**Fig. 3a**). Surprisingly, State 1 primarily composed of clusters 6 (highest signs of TCR activation; **Fig. 2e**) and 10. Cells in clusters 10, 1 and 9 showed a male-biased distribution within State 3, which is distinguishable by its stable *Tcf7* and gradually decreasing *Gzmb* levels (**Fig. 3b**). Clusters 2 and 7 were predominantly found in State 3 with no sex differences (**Fig. 3a**). Collectively, our analysis highlights the existence of CD8^+^ TIL differentiation from an activated *Tcf7*^−^ to a *Tcf7*^+^ state, with pertinent male bias occurring early in the process. To validate this finding, we used flow cytometry to investigate CD44^+^CD62L^−^CD8^+^ TILs before (day 7), during (days 9 and 10) and after (day 12) the sex-based bifurcation in MB49 growth (**Fig. 3c**). On day 7, most cells were Tcf1^high^Tim3^−^, which are reminiscent of primed T cells that have just migrated into the tumor. By days 9 and 10, Tcf1^high^Tim3^−^ cells rapidly converted to Tcf1^−^Tim3^+^ cells in both sexes, characteristic of cognate tumor antigen recognition and acquisition of an early effector program. On day 12, we observed gradual loss of surface Tim3 expression and a male-biased formation of Tcf1^low^Tim3^−^ cells, consistent with the trajectory that had been predicted from scRNA-seq data.

**Fig. 3.**
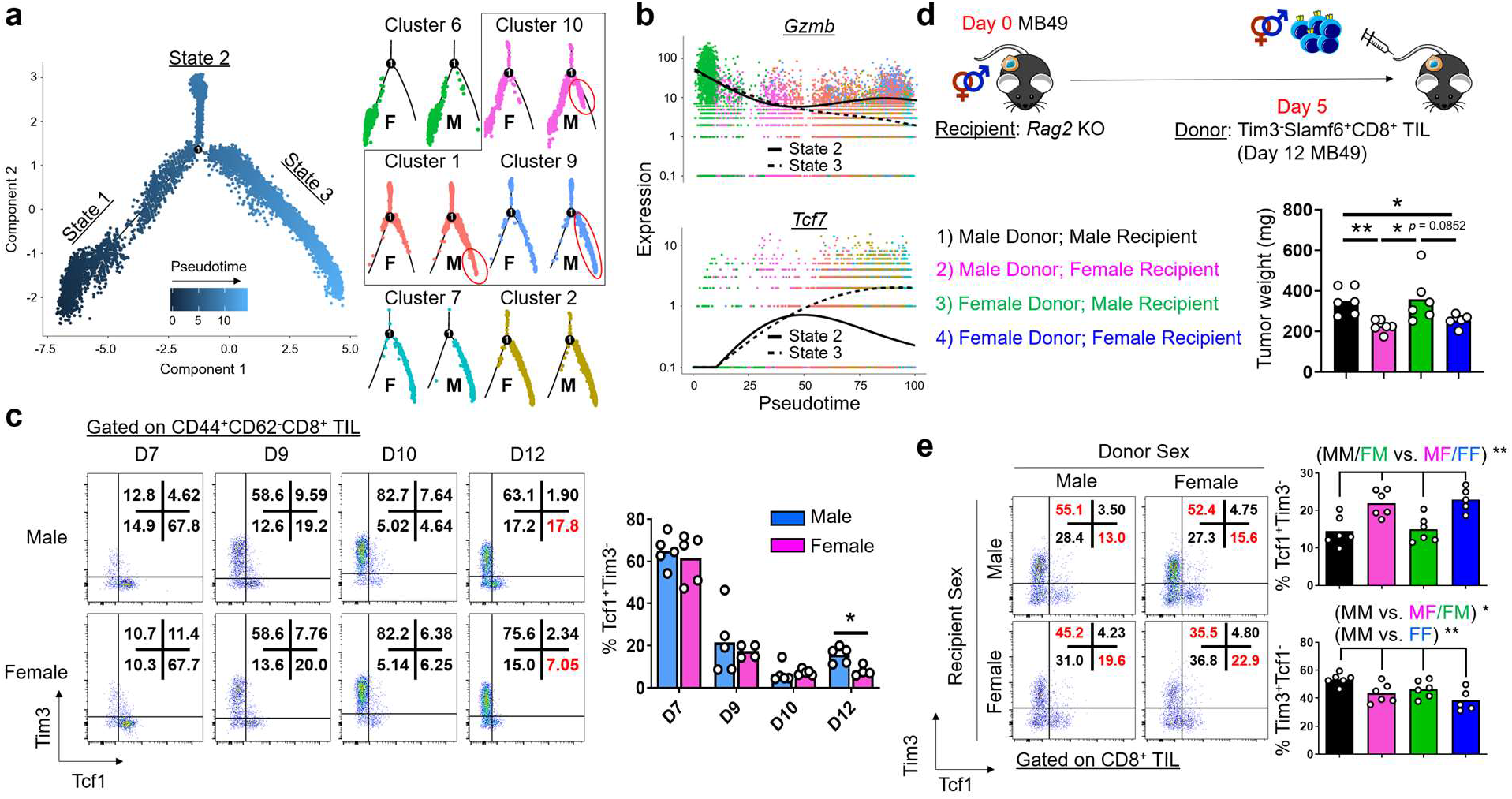
Male bias exists in CD8^+^ T cell commitment to exhaustion in the tumor microenvironment. **a**, Pseudotime analysis using Monocle-2^55^ across CD8^+^ T cells belonging to clusters 1, 2, 6, 7, 9 and 10 in day 10 MB49 tumors, from which 3 branches were identified: States 1 (progenitor), 2 and 3 (progeny). Each dot indicates a CD8^+^ T cell as colored by pseudotime (left; all cells) or cluster (right; stratified by sex). Boxed clusters show male-biased distribution in State 3 as illustrated in red. **b**, Expression of *Gzmb* and *Tcf7* across pseudotime for States 2 (solid) and 3 (dotted). **c**, Flow cytometric analysis of Tim3 and Tcf1 expression in CD44^+^CD62^−^CD8^+^ T cells from tumors of male and female mice 7, 9, 10 and 12 days post subcutaneous MB49 challenge. Left: Representative flow plots; Right: Frequency of Tcf1^+^Tim3^−^CD8^+^ T cells, with the box height representing the mean of all shown biological replicates. Blue: Male; Pink: Female. **d,** Growth of MB49 in *Rag2* KO mice that were adoptively transferred with 1.75 × 10^3^ Tim3^−^Slamf6^+^CD8^+^ T cells from day 12 MB49 tumors of WT mice. Colors for all possible donor-recipient combinations are listed. Tumor weights on day 14 are reported. **e,** Flow cytometric analysis of Tim3 and Tcf1 expression in donor TILs on day 14 from **d**. Left: Representative flow plots; Right: Frequency of Tcf1^+/−^Tim3^−/+^CD8^+^ T cells, with the box height representing the mean of all shown biological replicates. **c-e**, Statistical significance was determined by the Student’s *t* test. \**p* ≤ 0.05, \*\**p* ≤ 0.01.

An important question is if Tcf1^low^Tim3^−^CD8^+^ T cells can continue to differentiate within the tumor microenvironment. To address this question, we utilized an adoptive transfer model, in which we tracked the fate of these cells using Slamf6 as a cell surface surrogate for Tcf1 expression^11,24^ (**Fig. 3d**). Donor Slamf6^+^Tim3^−^CD8^+^ T cells gave rise to Tcf1^−^Tim3^+^ *bona fide* exhausted T cells that were unable to respond to re-stimulation (**Fig. 3e; Extended Data Fig. 4a**). As assessed by the lower frequency of Tcf1^low^Tim3^−^ and higher frequency of Tcf1^−^Tim3^+^ cells in male recipients, terminal differentiation of progenitor cells was most accelerated, resulting in the least tumor control, when the sex of both donor and recipient was male (**Fig. 3e**). Similarly, we observed male-biased accumulation of Tcf1^low^ progenitor exhausted and Tcf1^−^Tox^+^ exhausted CD8^+^ TILs in day 12 MB49 tumors of WT mice (**Extended Data Fig. 4b–4e**). These findings suggest that male bias goes beyond formation of progenitor exhausted CD8^+^ TILs and persists during their downstream differentiation. In humans, we observed a higher frequency of exhausted CD8^+^ TILs in men versus women when we probed basal cell carcinoma samples obtained prior to immunotherapy^32^ (“CD8_ex”; **Extended Data Fig. 5a and 5b**), as well as treatment-naïve non-small-cell lung cancer^33^ (“CD8_C6-LAYN”; **Extended Data Fig. 5e and 5f**). In addition, although CD8^+^ TILs in basal cell carcinoma primarily consisted of a memory-like phenotype (92% in women; 58% in men), they showed female bias in their expression of key effector molecules like *KLRG1* and *GZMB* (**Extended Data Fig. 5b**).

## Opposing transcriptional regulation of Tcf7 by androgen and type I IFN accompanies sex differences in CD8^+^ T cell early fate decision

MB49 growth in FCG mice varied by their sex organ rather than sex chromosome complement, suggesting that pertinent sex-specific differences in CD8^+^ T cell immunity are likely regulated by sex hormones (**Fig. 4a**). Upon analyzing scRNA-seq profiles of day 10 MB49 CD8^+^ TILs and lymphocytic choriomeningitis virus-specific CD8^+^ T cells^11^, we found that androgen, but not estrogen, signaling was uniquely enriched in clusters with expression profiles reflecting the Tcf1^low^ progenitor exhausted state (**Fig. 4b and 4d-4e; Supplementary Data 2**). In humans, androgen signaling also was enriched in activated, but less differentiated, CD8^+^ T cell clusters, while estrogen signaling showed variable enrichment (**Extended Data Fig. 5c and 5g**). Importantly, we identified a negative correlation between androgen and type I Interferon (IFN) signaling (Mouse: **Fig. 4c**; Human: **Extended Data Fig. 5d and 5h**), the latter of which has been shown to repress Tcf1 activity and inhibit the formation of progenitor exhausted CD8^+^ T cells^5,6^. We validated our scRNA-seq analyses with FACS-sorted CD8^+^ TILs at different stages of differentiation. Androgen receptor (*Ar*) mRNA not only showed a pan male bias, but also progressively increased – like *Tcf7* – from activated PD1^+^Slamf6^−^ to progenitor exhausted PD1^−^Slamf6^+^ populations (**Fig. 4f**). The PD1^+^Slamf6^+^ subset, representing an intermediate state that lacked markers of type I IFN exposure (i.e., *Isg15*), showed a male bias in *Tcf7* transcript levels (**Fig. 4f**).

**Fig. 4.**
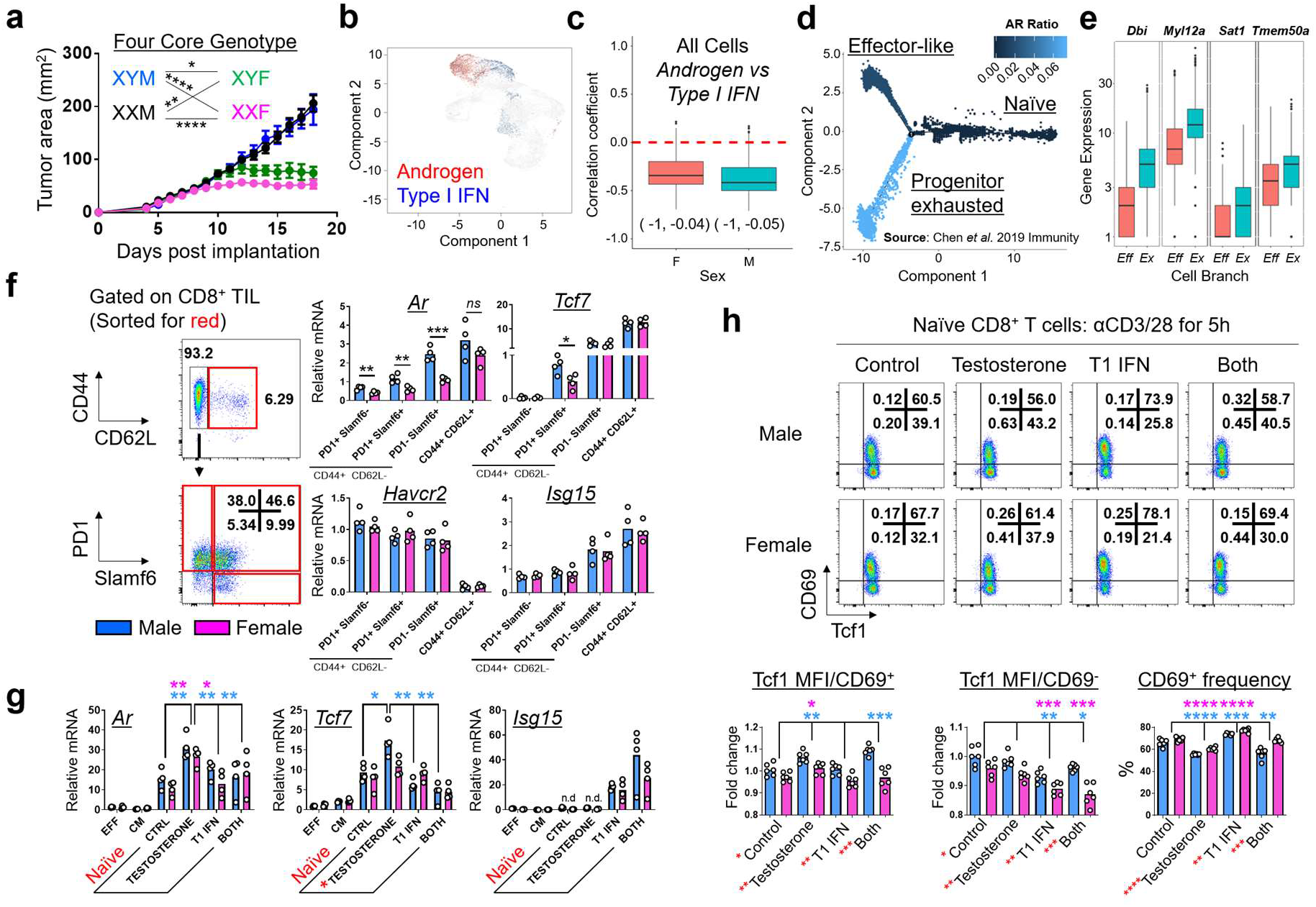
Opposing transcriptional regulation of *Tcf7* by androgen and type I IFN accompanies sex differences in CD8^+^ T cell early fate decision. **a**, Growth of MB49 in Four Core Genotype mice. Mean tumor area (mm^2^) ± SEM are reported, with statistical significance determined using the repeated measures two-way ANOVA. *n* = 7-11 mice per group. **b**, Enrichment of androgen response (red) and type I IFN (blue) signatures in CD8^+^ T cells from day 10 MB49 tumors. **c**, Correlation between androgen and type I IFN signatures, with 95% confidence intervals in brackets. **d,** Pseudotime analysis of virus-specific CD8^+^ T cells from Chen *et al* data^11^ using Monocle-2, as colored by the relative enrichment of androgen response signature. **e**, Boxplots show expression of genes from the androgen response signature in effector-like (“Eff”; orange) and progenitor exhausted (“Ex”; green) cells. **f and g**, qPCR analysis for indicated genes using **f)** FACS sorted CD8^+^ T cell subsets (red box) from day 14 MB49 tumors **g)** effector memory (“EFF”), central memory (“CM”) or naïve CD8^+^ T cells from spleens and peripheral lymph nodes. Naïve CD8^+^ T cells were stimulated with testosterone, type I IFN or both for 6 hours. Relative gene expressions are shown with the box height representing the mean of all shown biological replicates. **h,** Flow cytometric analysis of CD69 and Tcf1 expression in activated CD8^+^ T cells in the absence or presence of testosterone, type I IFN or both. Top: Representative flow plots; Bottom: Frequency of CD69^+^ cells and Tcf1 expression in CD69^+/−^ cells, with the box height representing the mean of all shown biological replicates. **f-h**, Statistical significance was determined by the Student’s *t*-test. \**p* ≤ 0.05, \*\**p* ≤ 0.01, \*\*\**p* ≤ 0.001, \*\*\*\**p* ≤ 0.0001. Blue: Male; Pink: Female. Red asterisks indicate significant sex differences.

Given these findings, we hypothesized that *Tcf7* undergoes opposing transcriptional regulation by androgen and type I IFN. Indeed, acute testosterone exposure significantly increased *Tcf7* in male CD8^+^ T cells, which was in turn blunted by type I IFN (**Fig. 4g**). Sex differences in the testosterone effect was likely not due to ligand recognition, as autoregulation of *Ar* was observed in both male and female T cells. We tested the consequences of testosterone exposure in male and female CD8^+^ T cells that were simultaneously activated *in vitro* through TCR/CD28 (**Fig. 4h**). Optimal T cell activation, as assessed by CD69 expression, was impaired in the presence of testosterone. While testosterone significantly increased Tcf1 expression in CD69^+^CD8^+^ T cells of both sexes, its effect was greater in magnitude and more resistant to type I IFN in males compared with females. Indeed, Tcf1 expression was higher in male CD69^+^CD8^+^ T cells even without acute hormonal treatment.

## Discussion

In contrast to prior studies that have largely focused on sex differences in tumor cell intrinsic biology in the settings of bladder^13,34^ and other cancers^35–40^, our work highlights the critical role of T cell immunity. We demonstrate that distinct CD8^+^ TIL fate determination underlies sex bias in spontaneous rejection of murine bladder tumor models, which provides a potential mechanism underlying sex differences in human bladder cancer incidence. Since male-biased predisposition for CD8^+^ T cell exhaustion appears as early as its commitment to the Tcf1^low^Tim3^−^ progenitor exhausted state that can be well-reinvigorated by anti-PD1 therapy^24–28^, our findings support the previously proposed notion that men may benefit more from immune checkpoint blockade^8,9^. On balance, molecular profiling efforts of clinical bladder cancer specimens suggest that basal subtype bladder cancers, which are more common in women^41^, may have increased therapeutic response to immune checkpoint blockade^34^.

Androgens are traditionally considered immunosuppressive^42,43^ and believed to contribute to male bias in bladder cancer risk in mice^12,44^ and humans^45^. Here, we show that androgens in the tumor microenvironment negatively impact T cell fate by regulating Tcf1/*Tcf7*. We also uncovered sex specificity in pertinent androgen effects of T cell exhaustion and type I IFN-mediated suppression, which may be due to a well-established female bias in the IFN response and downstream JAK/STAT signaling, both at baseline and upon active inflammation^7^ (**Extended Data Fig. 6**). This simultaneously sheds light on unresolved mechanisms by which type I IFN promotes CD8^+^ T cell terminal differentiation. As such, future work is necessary to disentangle the molecular circuitry of androgen, Tcf1 and type I IFN. In this regard, there exists significant precedent for the involvement of Nuclear Receptor family members, including *NR4A*^46–49^ and *NR3C1* (glucocorticoid receptor)^50^, in CD8^+^ T cell exhaustion. Finally, our work showing sex specific CD8^+^ TIL behavior in bladder cancer highlights the broader opportunities for discovery due to sex disparities in health and disease.

## Methods

### Mice

C57BL/6 (WT; Stock number 000664), C57BL/6 *Rag2*^−/−^ (008449), *Tcrb/Tcrd*^−/−^ (002122), *Ighm*^−/−^ (002288) and FCG^13^ (Four Core Genotype; 010905) mice were obtained from Jackson Labs. 5-12 weeks old mice, maintained in a specific pathogen-free environment, were used for experiments. Power analysis was not performed for sample size determination. Experiments – except for those utilizing FCG mice (Boston Children’s Hospital) – were conducted under protocols approved by the Institutional Animal Care and Use Committee at the Medical University of South Carolina and the Ohio State University.

### Tumor model

To induce bladder carcinogenesis, male and female WT and *Tcrb/Tcrd* knockout or FCG C57BL/6 mice were fed *ad libitum* with 0.1% BBN (TCI America) water for 14 weeks and then switched to normal water. All mice were monitored daily for morbidity (i.e. palpable tumor/abdominal swelling, hunched posture and urine staining around perineum). If mice survived the 40-weeks-long regimen, they were considered as censored from the Kaplan-Meier survival curve analysis. BKL171 was derived from the bladder tumor of an XXM FCG mouse at the end of a 40 weeks-long BBN regimen.

MB49 (a gift from C. Voelkel-Johnson from the Medical University of South Carolina) and BKL171 mouse urothelial carcinoma cells were cultured in Dulbecco’s modified Eagle’s medium with 10% heat-inactivated fetal bovine serum and 1% penicillin/streptomycin. 5 × 10^5^ tumor cells were resuspended in 100 μL ice cold PBS for subcutaneous injection into the right flank of a mouse. For antibody-mediated T cell depletion experiments, mice were injected intraperitoneally with 200 μg of anti-mouse CD4 (Clone GK1.5, BioXCell) and/or CD8 neutralizing antibodies (Clone 53-6.7, BioXCell), followed by 100 μg thereafter on the indicated days. Tumor surface area (width × length mm^2^) was measured using an electronic caliper every day starting on day 4 post implantation.

### Flow cytometry

Mouse spleens and tumors were mechanically disrupted, with the latter additionally subjected to digestion with 1 mg/mL Collagenase D (Roche) for 30 min at 37°C while shaken at 125 rpm. Excess volume of ice-cold PBS with 2% bovine serum albumin was used to inactivate the enzymatic activity. Red Blood Cell Lysis Buffer (BioLegend) was utilized on tissues before they were passed through 70 μm filters to prepare single cell suspensions. For cytokine production experiments, cells were re-stimulated by 50 ng/mL PMA (Sigma), 1 μg/mL Ionomycin (Sigma) and 1X Brefeldin A (BioLegend) in a 48-well plate for 2 hours at 37°C. For *in vitro* cultures, purified CD8^+^ T cells were left untreated or stimulated with 50 ng/mL testosterone (Sigma) and/or 50 U/mL mouse IFN alpha A (PBL Assay Science). Cells were stained in 4°C with eFluor506 fixable viability dye for 10 minutes (Invitrogen), followed by extracellular surface markers and FcR block concurrently for 30 minutes. All intracellular staining was performed using the Foxp3 transcription factor staining kit (Invitrogen) according to the manufacturer’s instructions. All samples were acquired on LSRFortessa or Cytek Aurora and analysis was performed using FlowJo VX or OMIQ, respectively.

Fluorochrome-conjugated antibodies directed against the following mouse antigens (Clone) were used: CD45 (30-F11); CD3 (AF-700/17A2), CD8 (53-6.7), CD4 (RM4-5), PD1 (J43), Tim3 (RMT3-23), Slamf6 (13G3-19D), Tox (REA473), Tcf1 (C63D9), CD44 (IM7), CD62L (MEL-14), CTLA4 (UC10-4B9), Lag3 (C9B7W), Klrg1 (2F1/KLRG1), T-bet (O4-46), Ki67 (SolA15), TNFα (MP6-XT22), IFNγ (XMG1.2), Granzyme B (12-8898-82) and CD69 (H1.2F3).

### Adoptive Transfer

CD8^+^ T cells were isolated by MACS (Miltenyi Biotec) from draining inguinal lymph nodes of MB49 bearing male and female mice 14 days post implantation (**Fig. 1c**). Alternatively, they were isolated by FACS from day 12 MB49 TILs in male and female mice based on Tim3^−^ Slamf6^+^ surface expression (**Fig. 3c**). Donor cells were intravenously administered into immunodeficient mice (*Tcrb/Tcrd* KO in **Fig. 1c**; *Rag2* KO in **Fig. 3c**). Recipient mice were injected with MB49 tumors at indicated time points and monitored for tumor growth.

### Single cell RNA sequencing

6 weeks old WT male and female C57BL/6 mice were inoculated subcutaneously with 5 × 10^5^ MB49 tumor cells. On day 10 post inoculation, single cell suspensions were prepared from the tumors after mechanical disruption and enzymatic digestion with 1 mg/mL Type IV Collagenase (Roche). Live tumor infiltrating CD8^+^ T cells (CD3^+^ CD8^+^ CD4^−^) were sorted on a BD FACSAria Ilu Cell Sorter and immediately processed for scRNA-seq. Experimental procedures for scRNA-seq followed established techniques using the Chromium Single Cell 3’ Library V3 Kit (10x Genomics). Briefly, FACS-sorted CD8^+^ T cells were loaded onto a 10X Genomics Chip A and emulsified with 3’ Single Cell GEM beads using a Chromium™ Controller. Libraries were constructed from the barcoded cDNAs (Translational Science Laboratory at the Medical University of South Carolina) and sequenced for approximately 300 million reads/sample on a NovaSeq S4 flow cell (Illumina) at the VANTAGE facility (Vanderbilt University Medical Center).

### Single-cell RNA-seq data analysis

Using the Cell Ranger software, we converted BCL files into FASTQ files, trimmed adapters and primer sequences, mapped reads to the mm10 reference genome, and quantified expression levels. In this step, to eliminate low-quality and dying cells, we filtered out cells with counts less than 200 and those with >5% mitochondrial counts. Then, we used the Seurat software^51^ for the downstream analysis, based on the count data obtained from Cell Ranger. Specifically, we normalized counts using the LogNormalize approach, visualized cells in a low-dimensional space using the Uniform Manifold Approximation and Projection (UMAP) algorithm^52^, and determined cell clusters using the shared nearest neighbor (SNN) modularity optimization based clustering algorithm^53^. This process resulted in identification of 11 cell clusters. Then, we identified cell type markers conserved between males and females for each cell cluster and also the genes that are differentially expressed (DE) between males and females using a Wilcoxon Rank Sum test and adjusted DE p-values for multiple testing using the Benjamini-Hochberg procedure^54^. For the pseudotime analysis, we used the Monocle 2 software^55^ with the count data for the cell clusters 1, 2, 6, 7, 9, and 10. Specifically, we ordered genes using the top 1000 genes with the largest variations in expression among these 6 cell clusters, reduced data dimensionality using the DDRTree algorithm^56^, and ordered genes along the trajectory. Gene set enrichment analyses were implemented using the hypergeometric test with the Kyoto Encyclopedia of Genes and Genomes (KEGG) gene sets obtained from the MSigDB (https://www.gsea-msigdb.org/gsea/msigdb/index.jsp).

### Secondary analysis of single-cell RNA-seq data

We implemented secondary analyses of single cell RNA-seq data obtained from previously published research. For Guo *et al.^33^*, we downloaded the count data from the GEO database with the accession number GSE99254 and selected only the cells corresponding to the CD8^+^ T cells from tumors. For Yost *et al.^32^*, we downloaded the count data for basal cell carcinoma from the GEO database with the accession number GSE123813, and we selected only the cells corresponding to CD8^+^ T cells and pre-treatment. Then, for both datasets, we used the Seurat analysis workflow described in “Single cell RNA-seq data analysis”. For Chen *et al.^11^*, we downloaded the count data from the GEO database with the accession number GSE131535. Then, we used the Monocle 2 analysis workflow described in “Single cell RNA-seq data analysis”.

### RNA Isolation and qPCR analysis

RNA was extracted from FACS/MACS-isolated CD8^+^ T cells and reverse-transcribed using RNeasy Micro Kit (Qiagen) and SuperScript™ IV VILO™ Master Mix with ezDNase™ Enzyme (ThermoFisher Scientific), respectively. Quantitative PCR was performed with the following primers:

*<u>Ar</u>*: forward, 5’-TCCAAGACCTATCGAGGAGCG-3’; reverse, 5’-GTGGGCTTGAGGAGAACCAT-3’;
*<u>Tcf7</u>*: forward, 5’-CCACTCTACGAACATTTCAGCA-3’; reverse, 5’-ACTGGGCCAGCTCACAGTA-3’;
*<u>Havcr2</u>*: forward, 5’-TCAGGTCTTACCCTCAACTGTG-3’; reverse, 5’-GGCATTCTTACCAACCTCAAACA-3’;
*<u>Isg15</u>*: forward, 5’-GGTGTCCGTGACTAACTCCAT-3’; reverse, 5’-CTGTACCACTAGCATCACTGTG-3’;
*<u>β-actin:</u>* forward, 5’-AGCTGAGAGGGAAATCGTGC-3; reverse, 5’-TCCAGGGAGGAAGAGGATGC-3’
*<u>Sry (Set 1):</u>* forward, 5’-TTGTCTAGAGAGCATGGAGGGCCATGTCAA-3’ reverse, 5’-CCACTCCTCTGTGACACTTTAGCCCTCCGA-3’
*<u>Sry (Set 2):</u>* forward, 5’-TGGGACTGGTGACAATTGTC-3’ reverse, 5’-GAGTACAGGTGTGCAGCTCT-3’

### Statistical analysis

Overall survival was analyzed using a log-rank test and tumor growth was analyzed by a two-way repeated measures ANOVA. Primary method of statistical analysis for other outcomes was a two-sided independent-sample *t*-test (Mann-Whitney U test in the event of non-normally distributed data). For all statistical testing, p-value < 0.05 was considered significant.

## Data availability

The datasets generated during and/or analysed during the current study are available from the corresponding author on reasonable request.

## Acknowledgment

We thank Cynthia Timmers and Marty Romeo from the Translational Science Laboratory at the Medical University of South Carolina for their assistance with the single cell RNA-sequencing efforts. We also thank Eugene Otlz for critical reading and editing of the manuscript. This work was supported by National Institutes of Health grants: P01 CA186866, R01 CA213290, R01 CA188419 and R01 AI077283 (Z. Li). H. Kwon was supported by Doctoral Foreign Study Award from Canadian Institutes of Health Research (201810DFS-422133-63414) and Graduate Fellowship from the Hollings Cancer Center in Charleston, SC, USA.

## Author contributions

H.K., X.L. and Z.L. conceived the project. H.K., D.C. and Z.L. designed experiments and wrote the manuscript. H.K. performed most experiments described herein and related analyses. H.K. and D.C. performed analyses of mouse and human scRNA-seq. S.K., A.L., L.Z., B.R. and NJ.S. contributed to *in vivo* tumour experiments. D.S., X.L and Z.L. provided intellectual input and critical edits to the manuscript. Z.L. supervised the project. All authors reviewed and approved the manuscript.

---

**Extended Data Fig. 1.**
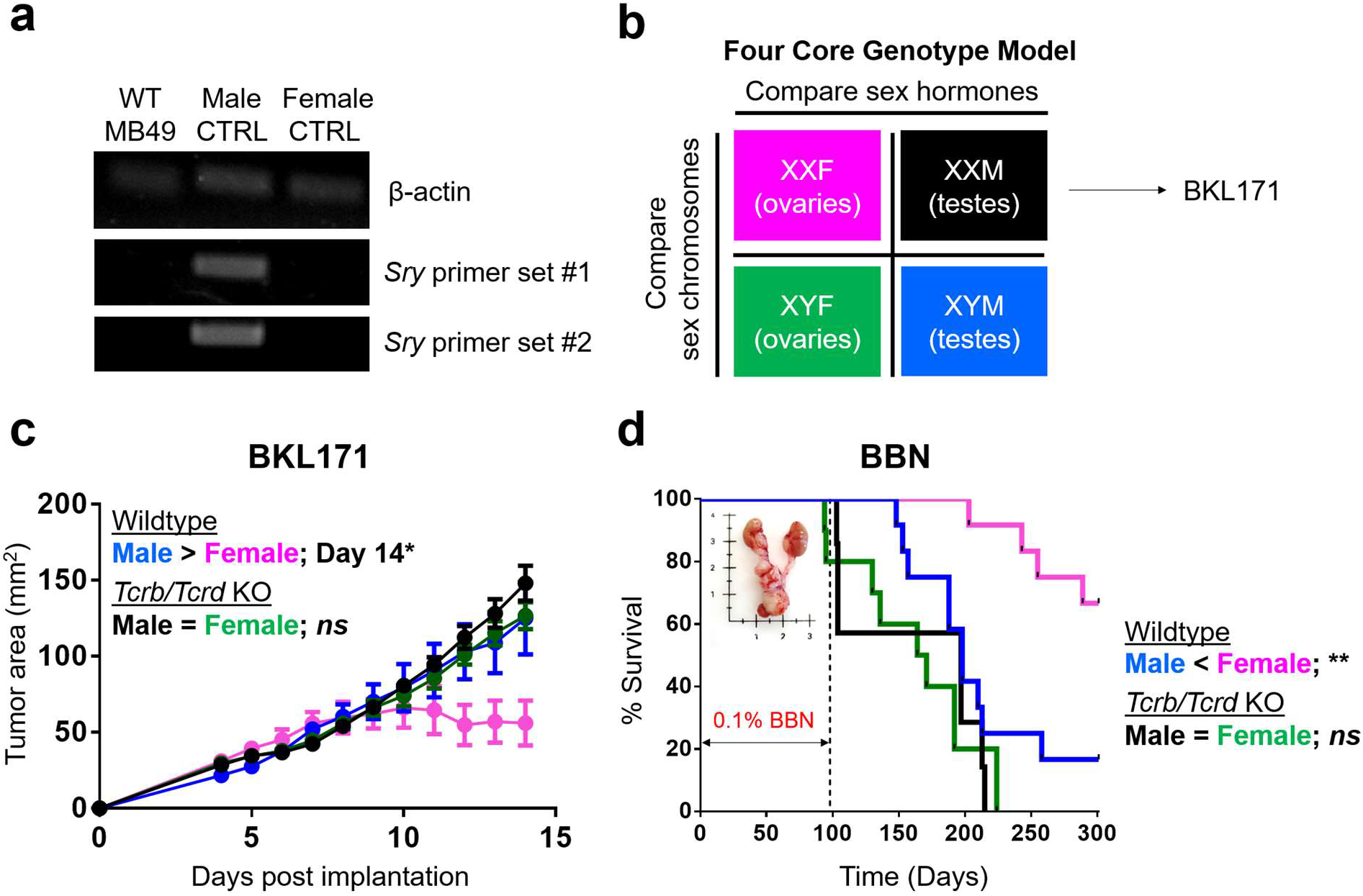
T cell immunity underlies sex biased outcomes in murine bladder cancer models. **a**, RT-PCR for qualitative detection of *β-actin* and Y-chromosome encoded *Sry* gene transcripts from MB49 cells. DNA extracted from tails of male and female mice are included as controls. **b**, Diagram representation of Four Core Genotype (FCG) mouse model. BKL171 was generated from BBN-induced bladder tumor of an XXM FCG mouse. **c**, Growth of BKL171 in mice with indicated genotypes after subcutaneous implantation. Mean tumor area (mm^2^) ± SEM are indicated, with following *p* values determined using the Student’s *t* test. Day 12 = 0.0753; Day 13 = 0.0776; Day 14 = 0.0487(*) for WT male and female (*n* = 4 each). No significant differences exist between *Tcrb/Tcrd* KO male and female (*n* = 10 and 9, respectively). **d**, BBN-induced carcinogenesis model. Mice are exposed *ad libitum* to 0.1% BBN in drinking water for the first 14 weeks to induce bladder cancer formation and then monitored for a total of 300 days. Percent survival is shown, with statistical significance determined using the log rank test. \*\**p* ≤ 0.01 between WT male and female (*n* = 12 each). No significant differences (*ns*) between *Tcrb/Tcrd* KO male and female (*n* = 7 and 10, respectively).

**Extended Data Fig. 2.**
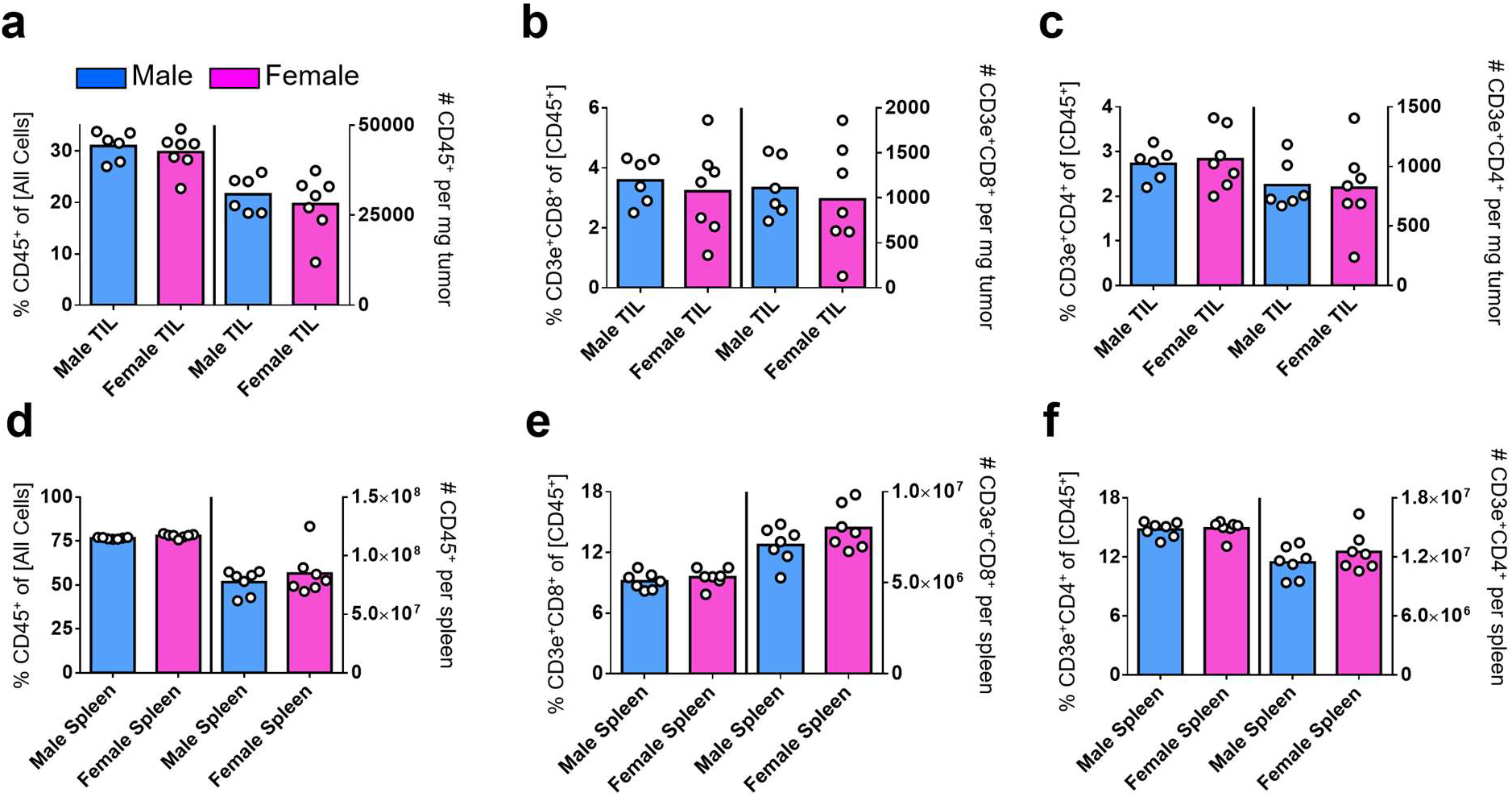
T cell numbers are comparable between male and female MB49 bearing mice. CD45^+^, CD3^+^CD8^+^ and CD3^+^CD4^+^ immune cell frequency and absolute number – as assessed by flow cytometry – in day 9 MB49 tumors (**a-c**) or spleens (**d-f**) are indicated in left and right y axes of each graph, respectively. Box height represents the mean of all shown biological replicates. No significant sex-based differences were detected. Blue = Male; Pink = Female.

**Extended Data Fig. 3.**
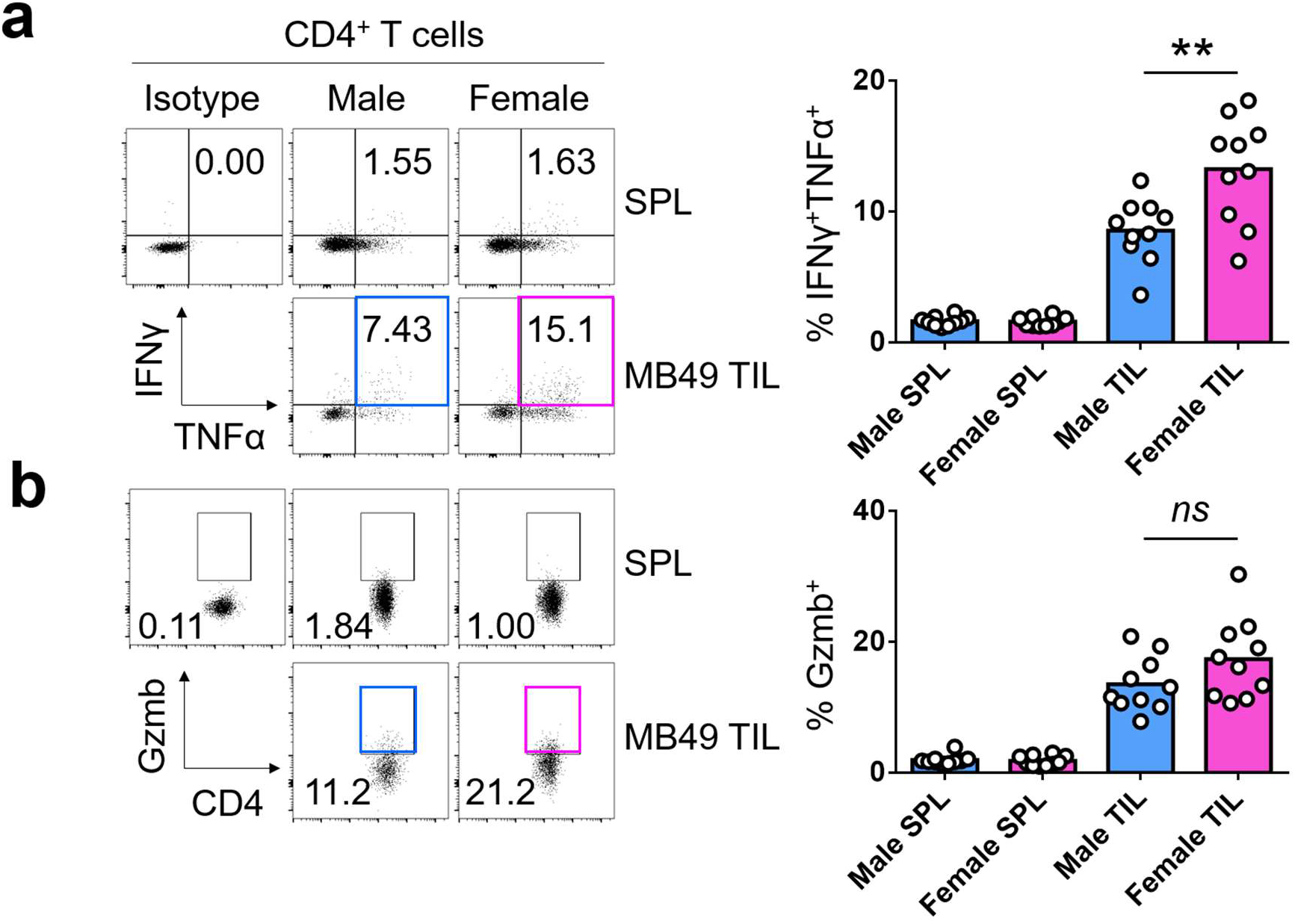
Female bias exists in CD4^+^ T cell effector response in the tumor microenvironment. Flow cytometric analysis of IFNγ and TNFα (**a**) and Gzmb (**b**) expression in CD4^+^ T cells from the spleens (SPL) and tumors (TIL) of male and female mice 9 days post subcutaneous MB49 challenge. Cells were stimulated *ex vivo* with 50 ng/mL PMA, 1 μg/mL Ionomycin and 1X Brefeldin A for 2 hours in 37 degrees Celsius. Left: Representative flow plots; Right: Frequency of CD8^+^ T cells expressing indicated effector molecules, with the box height representing the mean of all shown biological replicates. Statistical significance was determined using the Student’s *t*-test. \*\**p* ≤ 0.01. *ns* = not significant.

**Extended Data Fig. 4.**
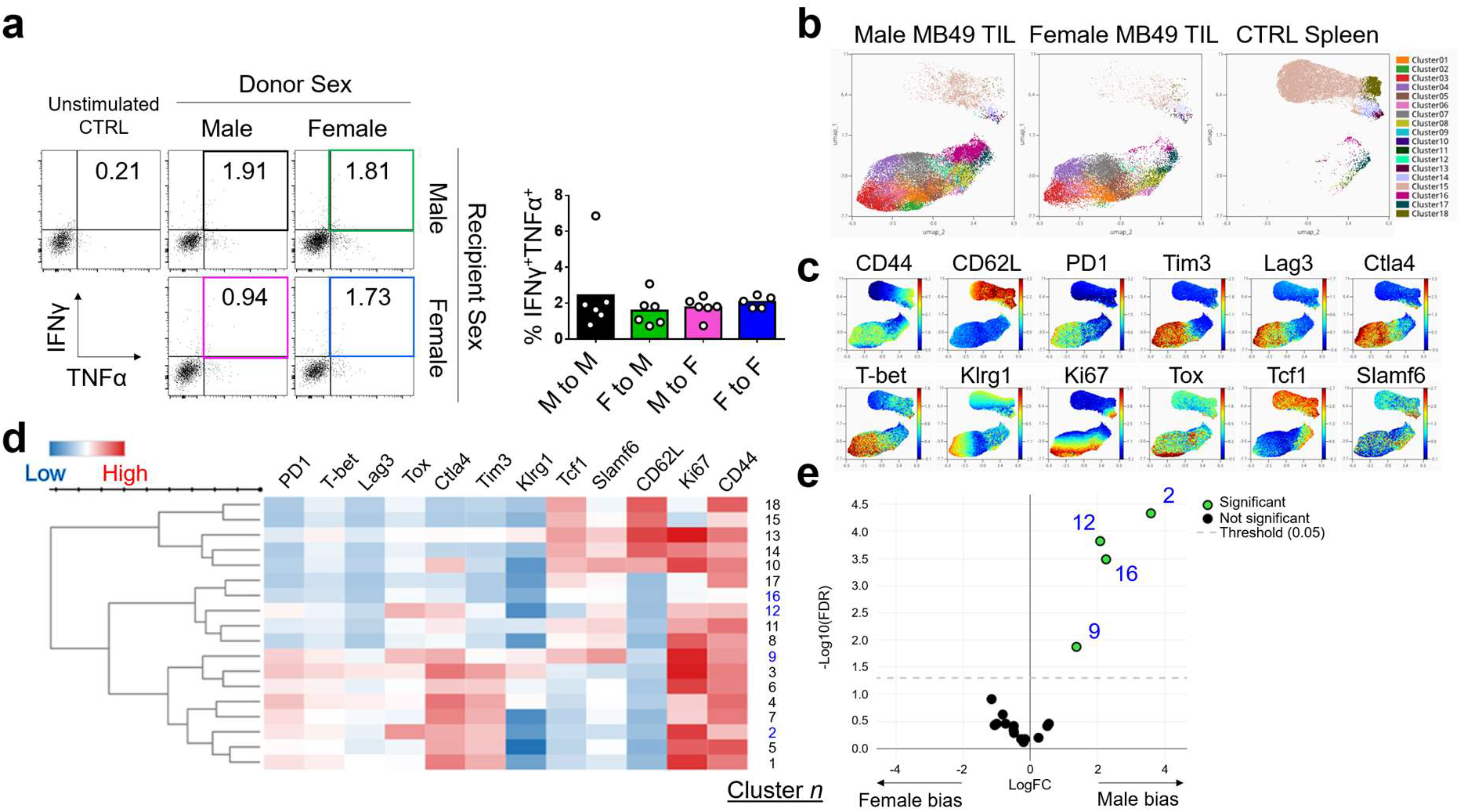
Male bias exists in CD8^+^ T cell commitment to exhaustion in the tumor microenvironment. **a**, Flow cytometric analysis of IFNγ and TNFα expression in donor male and female CD8^+^ T cells from tumors of male and female *Rag2* knockout recipient mice 12 days post subcutaneous MB49 challenge (see **Fig. 3d–e**). Cells were stimulated *ex vivo* with 50 ng/mL PMA, 1 μg/mL Ionomycin and 1X Brefeldin A for 2 hours. Left: Representative flow plots; Right: Frequency of CD8^+^ T cells expressing indicated effector molecules, with the box height representing the mean of all shown biological replicates. **b-e**, Spectral flow cytometry analysis of CD8^+^ T cells (Male = 22,875 cells; Female = 14,264 cells; Spleen control = 14,606 cells; *n* = 5 mice per sex) from day 12 MB49 tumors. **b**, UMAP as colored by cluster. **c**, Expression of indicated proteins in individual cells from **b**. **d**, Cluster visualization by heatmap. Clusters with differential abundance between sexes, as analyzed via EdgeR, are indicated by blue font and green circles in a volcano plot in **e**.

**Extended Data Fig. 5.**
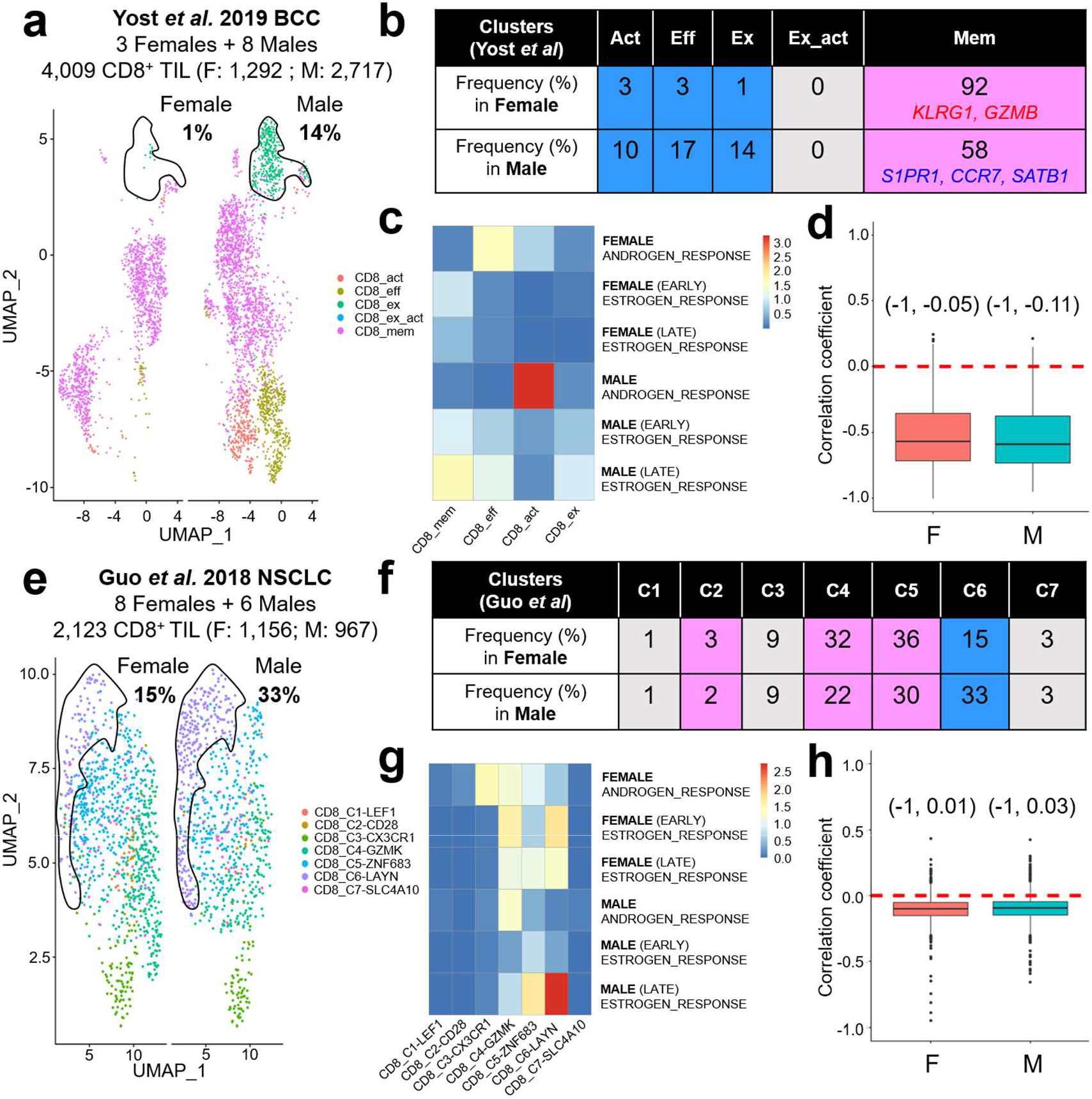
Male biased CD8^+^ T cell exhaustion in human cancers. Tumor infiltrating CD8^+^ T cells from basal cell carcinoma prior to immunotherapy^32^ (**a-d**) and treatment-naïve non-small cell lung cancer (**e-h**)^33^. T cell differentiation states were annotated the same way as published and stratified based on patients’ sex. Solid black lines enclose exhausted clusters that show male-biased frequency. **a,** Act = activated, Eff = Effector, Ex = Exhausted, Ex_act = Exhausted/activated, Mem = Memory. **e**, C1-LEF1 = naïve, [C2-CD28, C4-GZMK, C5-ZNF683] = intermediate between naïve and effector, C3-CX3CR1 = effector, C6-LAYN = exhausted, C7-SLC4A10 = mucosal associated invariant T (MAIT) cells. **b** and **f,** Frequency of indicated clusters in men and women. Blue = Male bias; Pink = Female bias; Gray = no bias. Genes listed under “Mem” show notable sex biased expression in memory CD8^+^ T cells. **c** and **g,** Heatmaps showing enrichment of indicated sex hormone signatures. Colors are based on p values in –log10 scale. **d** and **h,** Correlation between androgen response and type I Interferon signatures. Numbers indicate 95% confidence intervals.

**Extended Data Fig. 6.**
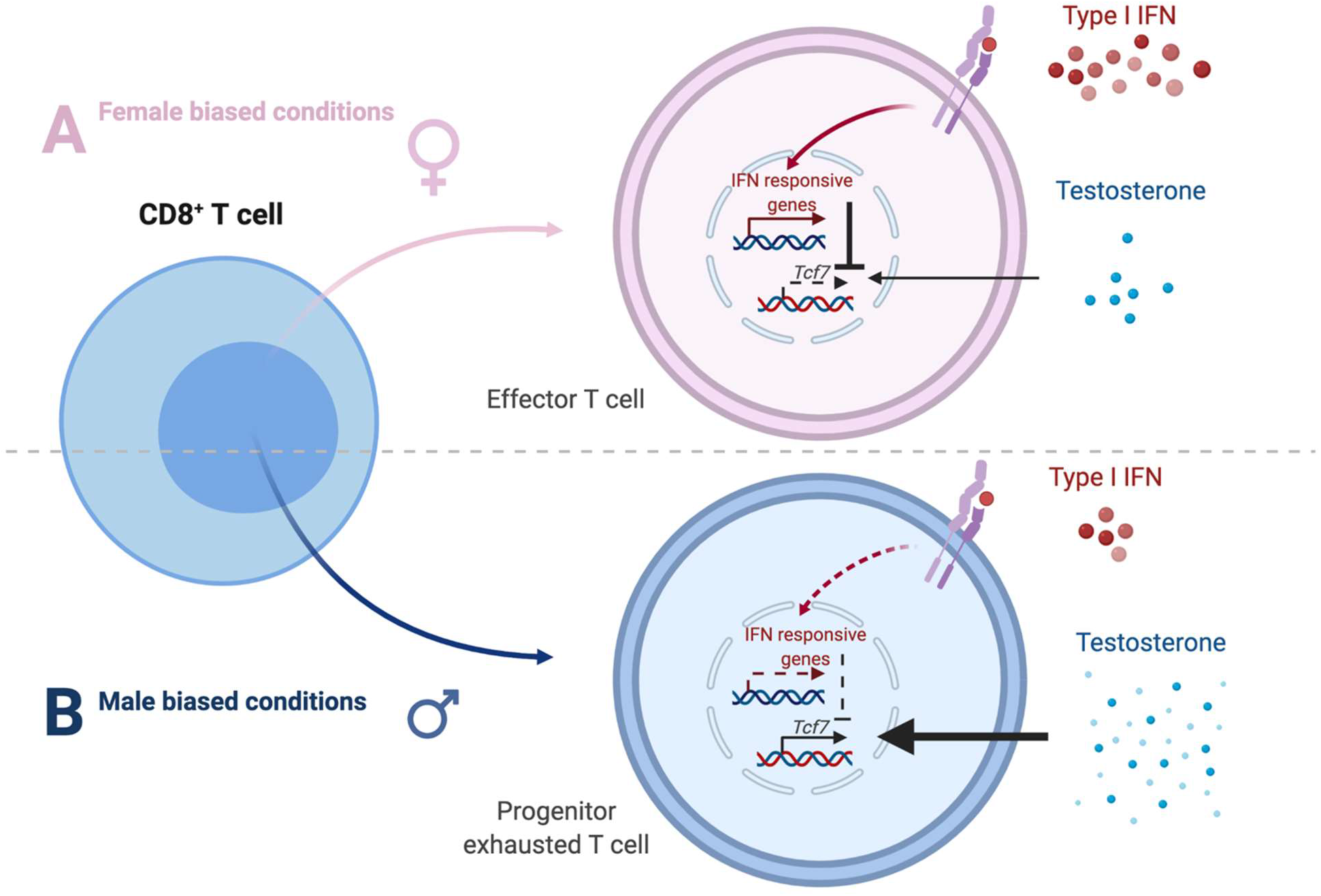
Schematic representation of sex differences CD8^+^ T cell fate.

**Supplementary Data 1 | Seurat_list_markers.xlsx**. Gene markers for 11 clusters of CD8^+^ T cells from day 10 MB49 tumors.

**Supplementary Data 2 | pvalue_hormone_IFN_by_sex.xlsx.** P values indicating the enrichment of the following signatures in 11 clusters of CD8^+^ T cells from day 10 MB49 tumors: 1) HALLMARK ANDROGEN RESPONSE, 2) HALLMARK ESTROGEN RESPONSE EARLY, 3) HALLMARK ESTROGEN RESPONSE LATE and 4) HALLMARK INTERFERON ALPHA RESPONSE.

